# On the hindlimb biomechanics of the avian take-off leap

**DOI:** 10.1101/2021.11.19.469279

**Authors:** E. A. Meilak, P. Provini, C. Palmer, N. J. Gostling, M. O. Heller

**Author notes:** Correspondence: Corresponding author, Prof. Dr.Markus O. Heller, University of Southampton, Faculty of Engineering and Physical Sciences, Department of Mechanical Engineering, Southampton, SO17 1BJ, United Kingdom.

## Abstract

Although extant land birds take to the air by leaping, generating the initial take-off velocity primarily from the hindlimbs, the detailed musculoskeletal mechanics remain largely unknown. We therefore simulated *in silico* the take-off leap of the zebra finch, *Taeniopygia guttata*, a model species of passerine, a class of bird which includes over half of all extant bird species. A 3D computational musculoskeletal model of the zebra finch hindlimb, comprising of 43 musculotendon units was developed and driven with previously published take-off ground reaction forces and kinematics. Using inverse dynamics, the external moments at the ankle, knee, and hip joints were calculated and contrasted to the cumulative muscle capability to balance these moments. Mean peak external flexion moments at the hip and ankle were 0.55 bodyweight times leg length (BWL) each whilst peak knee extension moments were about half that value (0.29 BWL). Muscles had the capacity to generate 146%, 230%, and 212 % of the mean peak external moments at the hip, knee, and ankle, respectively. Similarities in hindlimb morphology and external loading across passerine species suggest that the effective take-off strategy employed by the zebra finch may be shared across the passerine clade and therefore half of all birds.

## Introduction

Take-off is a crucial part of avian flight, requiring energy-intensive motion to accelerate into the air. Understanding how birds make the transition from standing statically to being airborne is one of the key components necessary for understanding how avian flight evolved. Previous work on the take–off of a variety of land birds demonstrated that the hindlimb plays a major role in propelling the bird in to the air, and suggests that the bipedal leap generates approximately 80-90% of the take-off velocity [1–6]. The group Passeriformes (passerines) includes over 5000 species of bird, making up approximately 60% of all bird species [7–9]. Among them, the zebra finch (*Taeniopygia guttata*), a frequently utilised model species, is also known to primarily use its hindlimbs to take to the air: previous studies showed that the hindlimb is responsible for producing 94% of their take-off velocity [2]. To date however, the detailed internal hindlimb mechanics necessary to produce a successful take-off remain largely unknown. This lack of knowledge in extant birds not only limits our understanding about how they master the feat of taking to the air but also presents an obstacle to accurately infer the capacity of fossil birds to be airborne, thus blurring our understanding of the origin of flight more generally.

Computational biomechanical modelling is a useful tool for calculating the internal mechanics occurring within a biomechanical system which are otherwise very difficult if not impossible to directly measure [10–15]. Moreover, the application of such computational analysis methods to extant animals is seen as a key strategy to systematically develop the sound biomechanical basis on which to further our understanding also of the conditions in extinct species [16–18]. The inverse analysis approach is one such computational method that takes measured kinematics and kinetics to drive a biomechanical model to calculate the internal moments at each degree of freedom of each joint [14, 19–22]. Detailed, 3D models of the musculoskeletal anatomy allow relating these external moments to the moment generating capacity of the internal force generating structures, i.e. the muscles, and to develop a more detailed understanding of internal avian hindlimb kinetics during the take-off leap. However, few studies have reported on the detailed external kinetics (ground reaction forces) [2, 4–6] associated with the avian take-off leap and even fewer studies have investigated 3D hindlimb kinematics for these activities [1]. The authors are aware of only a single study that has investigated the biomechanics of the avian jump, focussing on a predictive simulation of the ground dwelling elegant-crested tinamou *Eudromia elegans* [23]. However, due to the scarcity in ground reaction forces and kinematic data of the tinamou leap, data on which an inverse analysis would rely on, the authors opted for a forward approach to predict the leaping behaviour. In how far the simulations reflect kinetics and kinematics that are consistent with actual conditions that can be observed and measured in this bird therefore remains to be established. Although computational analyses in birds are available and have shown the value of such analyses to further our understanding of avian hindlimb biomechanics with respect to e.g. the critical role of the ankle muscles in the take-off leap of the tinamou and the function of the antitrochanter as a passive mechanism to stabilise the hip of the running ostrich [23, 24], to date no study has determined the internal hindlimb joint kinetics of a flying bird as it leaps into the air using detailed measurements of external forces and hind limb kinematics. With the application of X-ray reconstruction of moving morphology (XROMM) technology to capture detailed 3D bone kinematics of the avian take-off leap, in combination with computational biomechanical analyses, the technology is finally available to accurately simulate the internal mechanics of the extant avian take-off [1, 10, 16, 25–28].

The current study therefore combines external forces and detailed bone kinematics that feed into computer simulations into the biomechanics of the hindlimb throughout the take-off leap of the zebra finch. In doing so we aim to address the following hypotheses that help to develop a comprehensive understanding of the mechanical requirements birds need to meet to propel themselves into the air. We firstly hypothesised that in order to generate the motion of the avian take-off leap, characterised by the hip, knee, and ankle joints all extending until the bird is airborne [1, 3], net external flexion moments of similar peak magnitudes act at all these joints which the muscles balance by exerting extension moments. Consistent with the observation that predicative simulations of the tinamou predict the ankle to be most critical in determining the success of the take-off leap in a ground-dwelling bird [23], we further hypothesised that the ankle extensors of the zebra finch possess the largest capacity to balance the external moments. Finally, considering the avian hips’ powerful capacity to generate internal/external rotation (IER) moments and a poor capacity to generate abduction/adduction (ABAD) moments [10, 16, 24] in the presence of the antitrochanter, we hypothesise that external moments of similar peak magnitudes act in IER/ABAD on the hip joint of the zebra finch and that the bird possess powerful ability to actively balance the IER moments.

## Materials and methods

To address our hypotheses, we built upon previously published detailed kinetics and kinematics data of the take-off leap of the zebra finch as input to a biomechanical simulation of the avian take-off leap [1, 2] as explained in more detail below.

### Overview

Key steps of our analysis methodology included the use of CT scans of the same individual, together with additional morphological data in the literature to characterise the bone and musculature respectively, from which a detailed 3D musculoskeletal model of the zebra finch hindlimb was developed. Here, muscles were mapped from a previously published magpie musculoskeletal model on to the zebra finch skeleton [10, 29, 30]. In addition, previously published 3D kinematics and ground reaction forces were used to drive the musculoskeletal model in inverse dynamics analyses to estimate external joint moments [1, 2]. The combination of these unique datasets, collected from the same species and even the same individual, offered a unique opportunity to generate an accurate simulation of the zebra finch take-off biomechanics.

The external joint moments were compared against the moment generating capacity of the muscles to document the biomechanical requirements and to assess the zebra finch’s capability to actively balance the hindlimb joint moments experienced throughout the take-off leap. Here, moments about hip flex/extension (FE), ab/adduction (ABAD), int/external rotation (IER) and moments about FE at both the knee and ankle joints were considered. In addition, by comparing the hindlimb morphology and ground reaction forces of a variety of passerines ranging in mass (zebra finch *Taeniopygia guttata*, 15.4 g, starling *Sturnis vulgaris*, (77.3 g), crow *Corvus corone* (440 g) and raven *Corvus corax*, 1.1 kg) it was possible to explore the take-off mechanics of passerines more generally.

### Materials, model building, and musculoskeletal analysis approach

The computational biomechanical musculoskeletal model of a zebra finch (15.4 g) was developed based on CT scans and muscle data of corvids from previously published works [1, 10, 29]. The CT data was used for establishing the 3D skeletal model. In order to describe the spatial relationships between bones, local coordinate systems were established based on shape fitting techniques and functional analyses of the joints [31], as detailed in the supplementary information (S1A). Muscles were modelled by 3D lines of action [12, 32, 33] by mapping muscles from the magpie skeleton to the zebra finch using elastic registration [34], while muscle maximum isometric force was scaled by body mass. Detailed kinematics derived from previously published XROMM data [1] were then used together with ground reaction forces [2] of zebra finch take-off leaps to drive the model in inverse dynamics simulations to calculate the external moments acting at the hindlimb joints. Nine sets of ground reaction forces obtained from 9 jumps of 4 birds (15.4±1.8g), were temporally synchronised with two sets of kinematics trials (taken from 1 bird with a mass of 15.4 g), resulting in 18 simulations analysed. Muscle moment arms were measured and used to calculate the maximum moment generating capacity about hip flexion/extension, ab/adduction and internal/external rotation, and knee and ankle flexion/extension. The muscle moment generating capacity was compared to the external joint moment to ascertain the zebra finch’s ability to balance the external joint moments.

### Skeletal model

The skeletal model was derived from CT scans (isotropic resolution 0.04mm) of a zebra finch *Taeniopygia guttata* specimen (15.4 g), obtained in previously published studies (for full details please refer to [1]). For the current study, bones were segmented using ITK-SNAP [35], and imported in to Rhino (v7; Robert McNeel & Associates, Seattle, USA) [36] where triangulated bone surfaces were obtained after fitting subdivision surfaces using the QuadRemesh function. Bones which were treated in this way were the pelvis, femur, patella, tibiotarsus and tarsometatarsus. The detailed definition of joint centres and axes and local bone coordinate systems are available in the supplementary information (S3).

A linked rigid body model with 4 segments including the pelvis, thigh, shank and tarsometatarsus was defined in OpenSim v4.1 [37]. Here, body segments were linked by 3 joints (hip, knee and ankle joints) with 3 rotational degrees of freedom (DoF) at the hip (allowing flexion/extension, internal/external rotation and ab/adduction) and 2 DoF at each of the knee and ankle joints (allowing flexion/extension and internal/external rotation). Because the foot remained stationary on the perch and motion of the trunk was largely due to the extension of the hip, knee and ankle joints, the foot was not included as a dedicated structure of the musculoskeletal model. In order to better capture the mechanics of hindlimb extension, a biomechanical model of the patella-femoral joint was incorporated in to the model. Here, the motion of the patella was defined by a 3D spline curve following approximately the trochlear surface of the femur. The patella was allowed to move along that spline as a function of the knee angle, with details of the motion informed by a musculoskeletal model of the helmeted guineafowl *Numida meleagris* [38]. In order to map patellar motion from the guineafowl to the zebra finch model, the femoral surface of the guineafowl model was elastically registered to the zebra finch femur [39]. The parameters of the elastic registration were then used to map the spline defining patellofemoral motion from the guineafowl to the zebra finch model using the R-package MesheR [34].

### Muscle geometry

The musculoskeletal model of the zebra finch included 43 muscles crossing the hip, knee and ankle, which were modelled as 3D polylines spanning origin and insertion using via points to fully describe their curved paths (Figure 1). Magpies (*Pica pica*) and zebra finch, share broadly similar hindlimb myology both following the characteristic hindlimb morphology of passerine birds [9]. Muscle attachment sites and via points were therefore mapped from a previously established model of the magpie [10] to the zebra finch hindlimb. Using a non-rigid iterative closest point (ICP) registration [39] the magpie femur, tibiotarsus and tarsometatarsus were elastically registered to the corresponding long bones of the zebra finch hindlimb [34] (Figure 2). Using 3D Slicer 4.11 [40], the pelvis of the magpie and zebra finch were first split in to the ilium, ischium and pubis before elastically registering them onto each other following the same approach as described for the long bones above. Following non-rigid registration, the rigid transformation and isotropic scaling parameters were recovered using Ordinary Procrustes Analysis (OPA) [41] computed between the vertices of the original magpie bone surface and the vertices of the magpie bone surface that was elastically registered to the respective zebra finch bone (Matlab (2019b, The Mathworks, Nantucket, USA). This step was encoded in a 4×4 homogeneous transformation matrix. The remaining difference between the positions of the OPA mapped vertices and their elastically registered counterparts was captured in a dense deformation field. The homogeneous transformation matrix and the dense deformation field were then applied to all attachment and via points of the magpie muscles associated to the respective bone surface using the R-package MesheR [34], and in so doing mapping muscles from the magpie hindlimb to the zebra finch (Figure 2). Wrapping cylinders and spheres were added with positions, orientations and radii individually adjusted to define the 3D muscle paths throughout the jumping RoM that avoided any intersection of muscles with bones.

**Figure 1.**
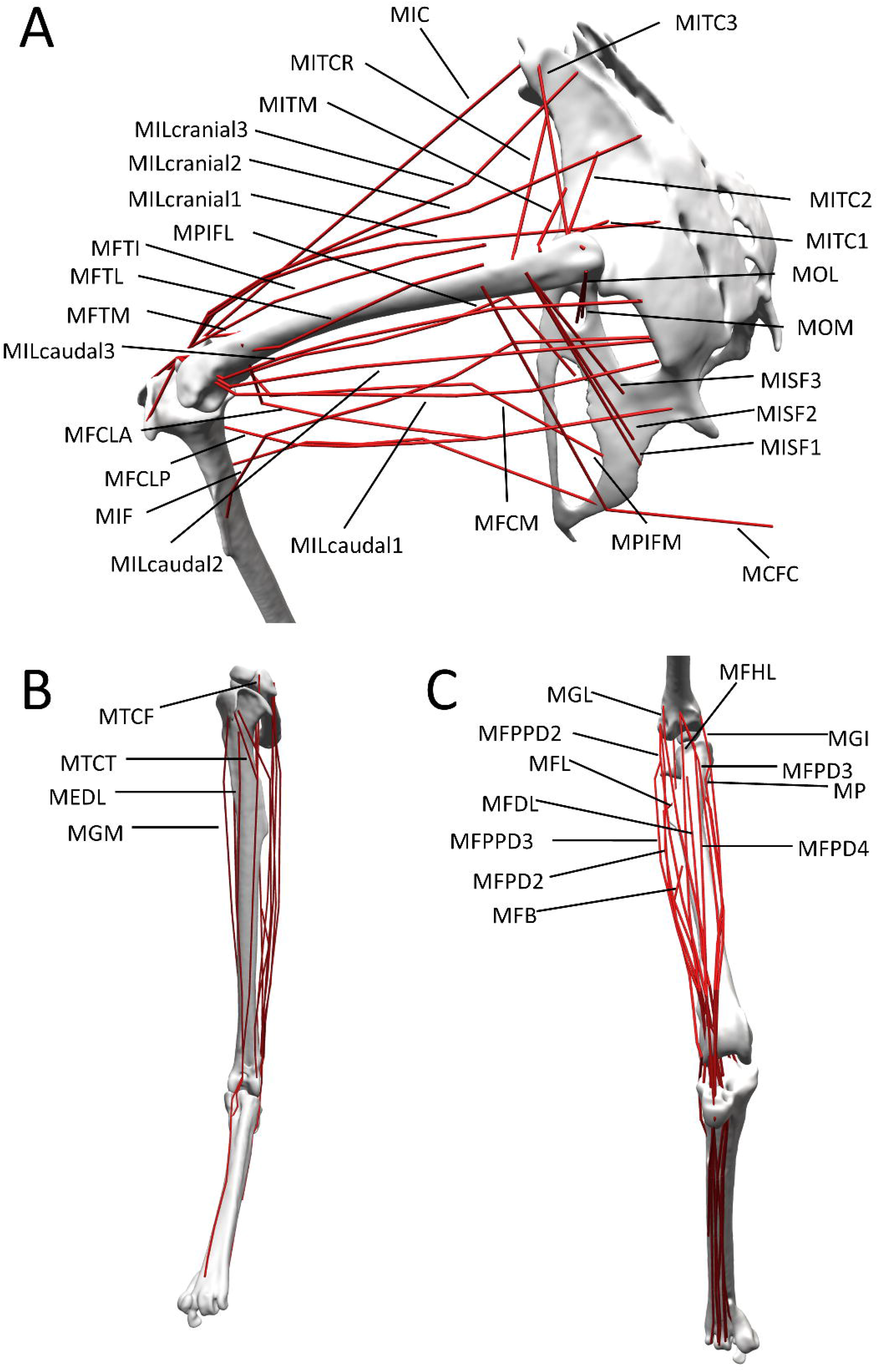
The musculoskeletal model of the left zebra finch hindlimb included 43 musculotendon units. A) Muscles that either cross the hip or knee joint and biarticular muscles crossing both hip and knee joints are shown in a caudo-lateral view. B) Anterior and C) posterior view of muscles crossing the ankle joint and biarticular muscles crossing both knee and ankle joints. For an explanation of the abbreviations of the muscles used here, please refer to Table 1.

**Table 1.**
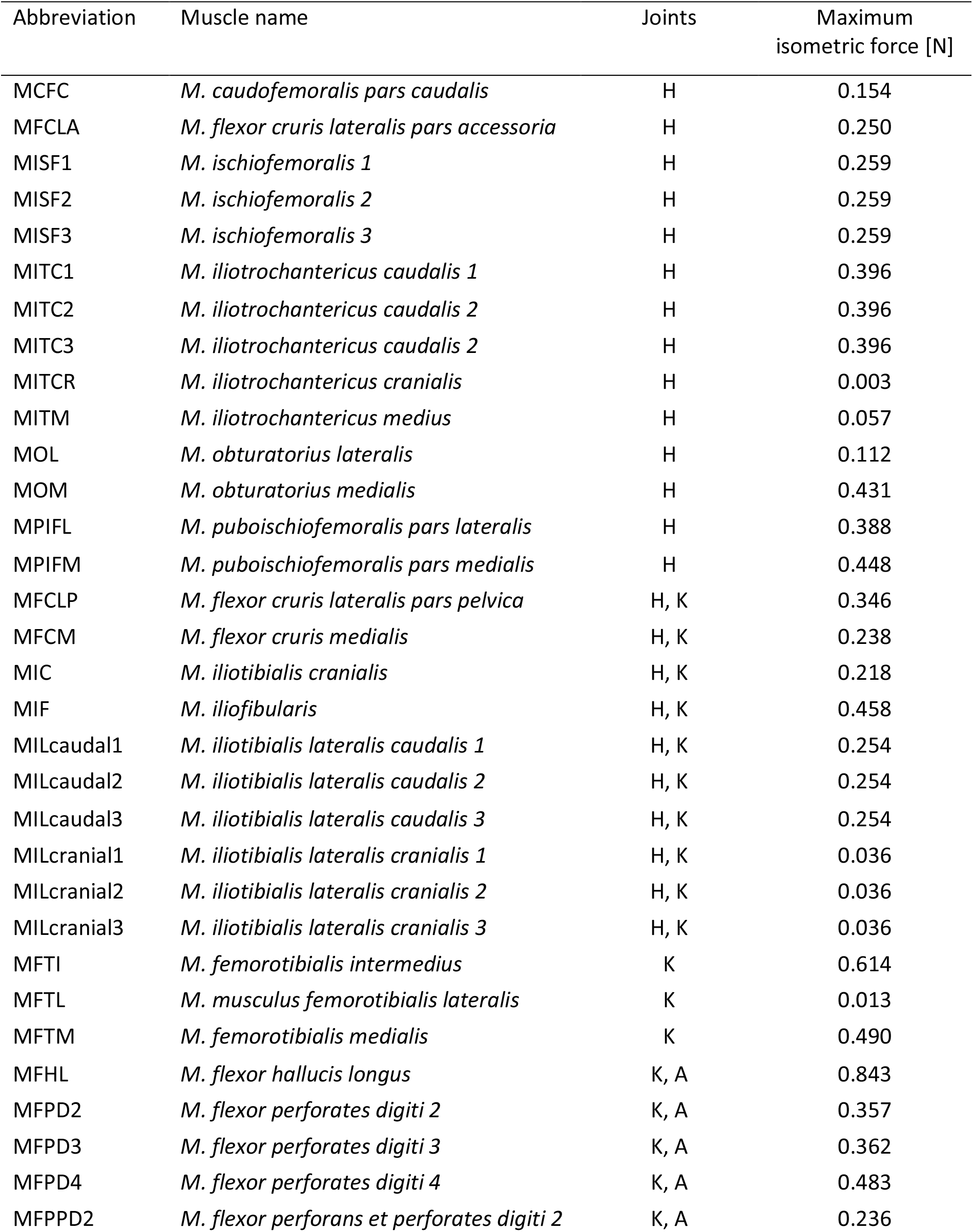
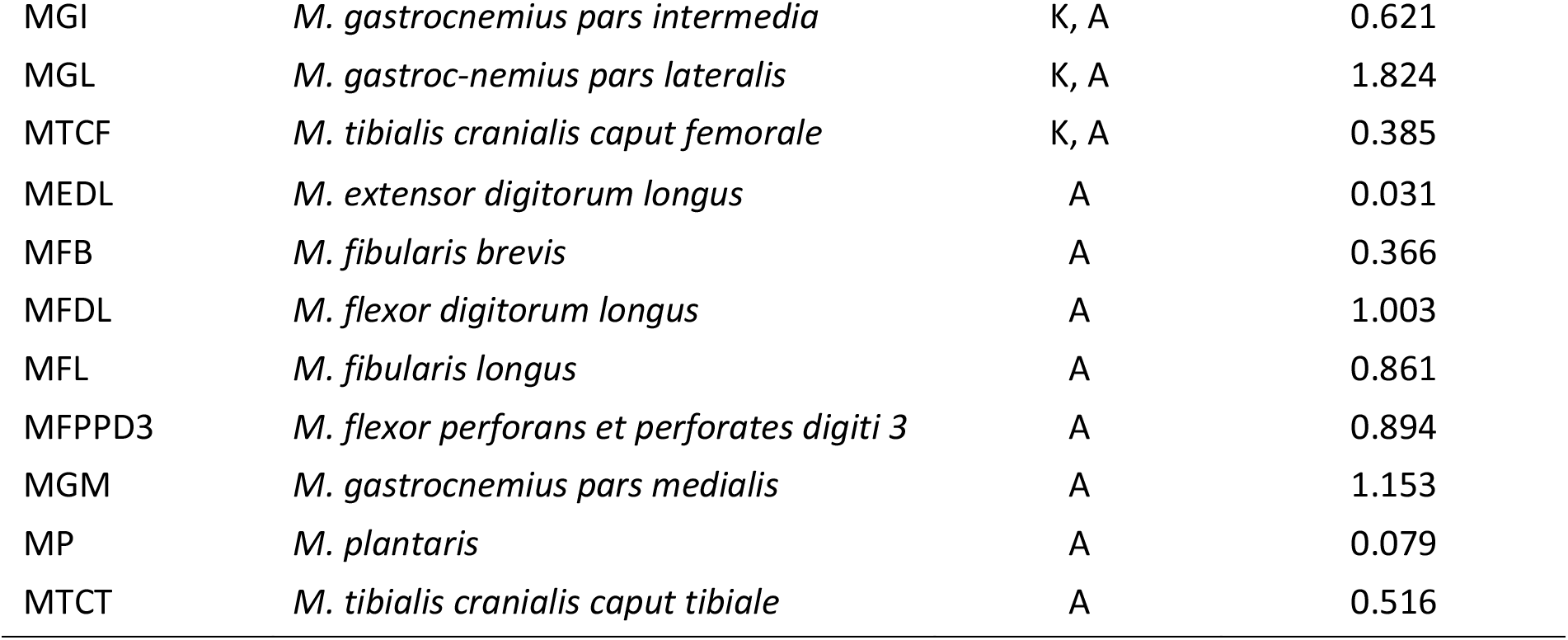
Musculotendon units included in the musculoskeletal model of the zebra finch, grouped by which joints they cross; H, K and A denote the hip, knee and ankle joints, respectively. Muscles which are categorised under two joints are biarticular. Maximum isometric force was calculated by scaling the maximum isometric force of the corresponding muscles of the magpie by mass [10].

| Abbreviation | Muscle name | Joints | Maximum isometric force [N] |
| --- | --- | --- | --- |
| MCFC | *M. caudofemoralis pars caudalis* | H | 0.154 |
| MFCLA | *M. flexor cruris lateralis pars accessoria* | H | 0.250 |
| MISF1 | *M. ischiofemoralis 1* | H | 0.259 |
| MISF2 | *M. ischiofemoralis 2* | H | 0.259 |
| MISF3 | *M. ischiofemoralis 3* | H | 0.259 |
| MITC1 | *M. iliotrochantericus caudalis 1* | H | 0.396 |
| MITC2 | *M. iliotrochantericus caudalis 2* | H | 0.396 |
| MITC3 | *M. iliotrochantericus caudalis 2* | H | 0.396 |
| MITCR | *M. iliotrochantericus cranialis* | H | 0.003 |
| MITM | *M. iliotrochantericus medius* | H | 0.057 |
| MOL | *M. obturatorius lateralis* | H | 0.112 |
| MOM | *M. obturatorius medialis* | H | 0.431 |
| MPIFL | *M. puboischiofemoralis pars lateralis* | H | 0.388 |
| MPIFM | *M. puboischiofemoralis pars medialis* | H | 0.448 |
| MFCLP | *M. flexor cruris lateralis pars pelvica* | H, K | 0.346 |
| MFCM | *M. flexor cruris medialis* | H, K | 0.238 |
| MIC | *M. iliotibialis cranialis* | H, K | 0.218 |
| MIF | *M. iliofibularis* | H, K | 0.458 |
| MILcaudal1 | *M. iliotibialis lateralis caudalis 1* | H, K | 0.254 |
| MILcaudal2 | *M. iliotibialis lateralis caudalis 2* | H, K | 0.254 |
| MILcaudal3 | *M. iliotibialis lateralis caudalis 3* | H, K | 0.254 |
| MILcranial1 | *M. iliotibialis lateralis cranialis 1* | H, K | 0.036 |
| MILcranial2 | *M. iliotibialis lateralis cranialis 2* | H, K | 0.036 |
| MILcranial3 | *M. iliotibialis lateralis cranialis 3* | H, K | 0.036 |
| MFTI | *M. femorotibialis intermedius* | K | 0.614 |
| MFTL | *M. musculus femorotibialis lateralis* | K | 0.013 |
| MFTM | *M. femorotibialis medialis* | K | 0.490 |
| MFHL | *M. flexor hallucis longus* | K, A | 0.843 |
| MFPD2 | *M. flexor perforates digiti 2* | K, A | 0.357 |
| MFPD3 | *M. flexor perforates digiti 3* | K, A | 0.362 |
| MFPD4 | *M. flexor perforates digiti 4* | K, A | 0.483 |
| MFPPD2 | *M. flexor perforans et perforates digiti 2* | K, A | 0.236 |
| MGI | *M. gastrocnemius pars intermedia* | K, A | 0.621 |
| MGL | *M. gastroc-nemius pars lateralis* | K, A | 1.824 |
| MTCF | *M. tibialis cranialis caput femorale* | K, A | 0.385 |
| MEDL | *M. extensor digitorum longus* | A | 0.031 |
| MFB | *M. fibularis brevis* | A | 0.366 |
| MFDL | *M. flexor digitorum longus* | A | 1.003 |
| MFL | *M. fibularis longus* | A | 0.861 |
| MFPPD3 | *M. flexor perforans et perforates digiti 3* | A | 0.894 |
| MGM | *M. gastrocnemius pars medialis* | A | 1.153 |
| MP | *M. plantaris* | A | 0.079 |
| MTCT | *M. tibialis cranialis caput tibiale* | A | 0.516 |

**Figure 2.**
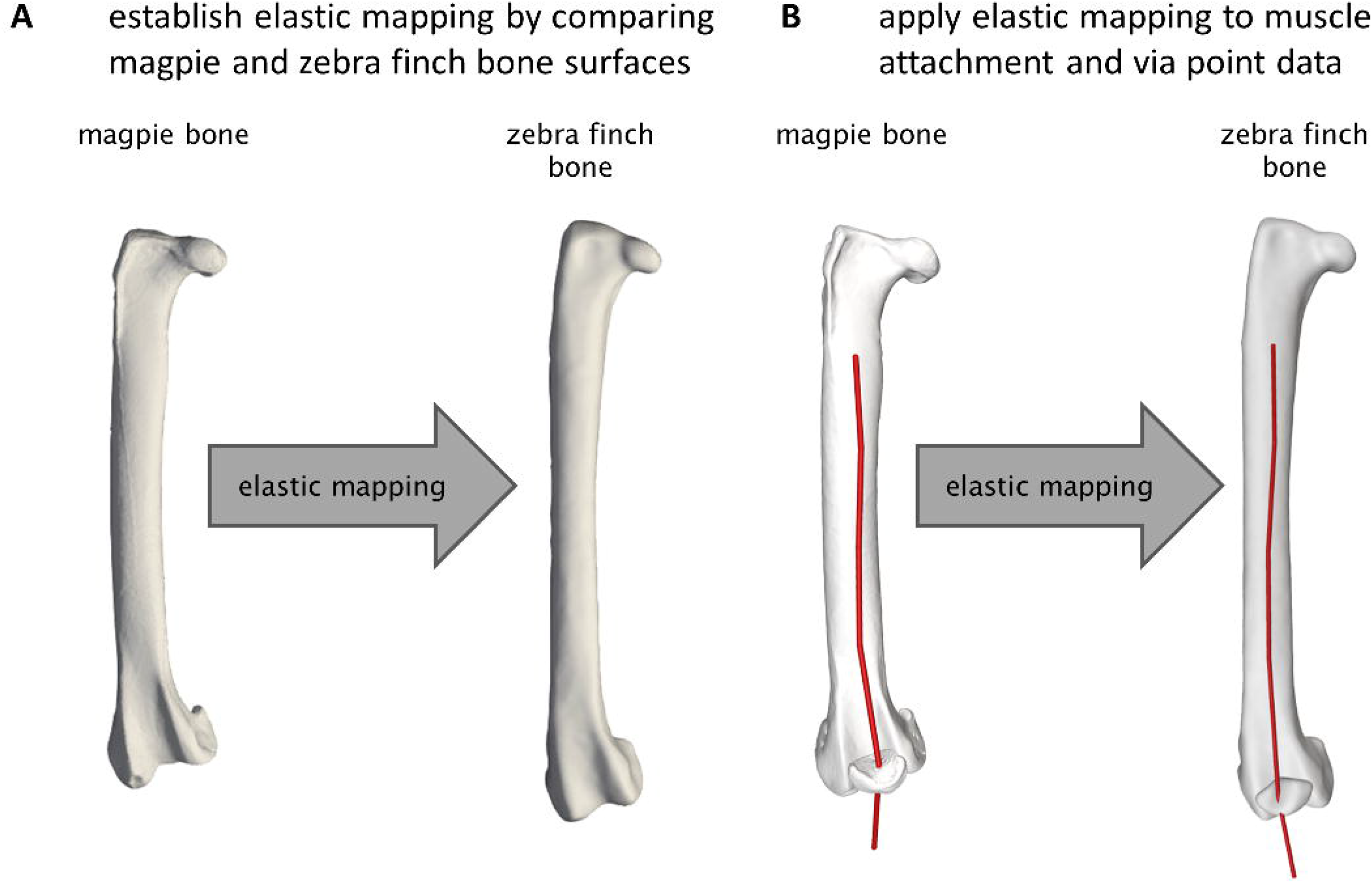
Schematic demonstrating the process of morphing the musculature from the magpie to the zebra finch hindlimb. To allow for a better visual comparison of the bones which differ by almost a factor of two in size, the bone surfaces depicted here are isotopically scaled by the reciprocal of the square root of their respective surface area. **A** Establishing an elastic mapping by comparing magpie and zebra finch bone surfaces. Here, magpie bones were first registered to the corresponding zebra finch bone using non-rigid iterative closest point (ICP) registration [39]. Following non-rigid registration, the rigid transformation and isotropic scaling parameters were recovered using Ordinary Procrustes Analysis (OPA) computed between the vertices of the original magpie bone surface and the vertices of the magpie bone surface that was elastically registered to the respective zebra finch bone. This step was encoded in a 4×4 homogeneous transformation matrix. The remaining difference between the positions of the OPA mapped vertices and their elastically registered counterparts was captured in a dense deformation field. **B** Application of the elastic mapping to muscle attachment and via point data. The homogeneous transformation matrix and the dense deformation field were then applied to all attachment and via points of the magpie muscles associated to the respective bone surface, resulting in an elastic registration of these structures to the zebra finch model.

### Biomechanical analysis

#### Kinematics

Detailed 3D kinematics were derived from previously published X-ray reconstruction of moving morphology (XROMM) data of the left hindlimb throughout two autonomous take-off leaps of the zebra finch [1]. For these analyses three tantalum bead markers with an approximate diameter of 0.5 mm were attached to the tarsometatarsus, while two markers were implanted to the tibia, one to the femur, and one at the pelvis. In order to reliably track 3D skeletal kinematics for the current study, the location of the pelvis was determined using the implanted pelvic marker and, while its orientation was determined using scientific rotoscoping, a reliable process in particular for bones with specific shapes such as the pelvis [42]. Owing to the generally higher precision of the 3D marker positions compared to 3D position and orientation data derived from scientific rotoscoping [1, 42], a method for reconstructing 3D skeletal motion that maximised usage of the marker data while minimising reliance on the manual process of scientific rotoscoping was developed. Here, 3D positions and orientations of the long bones of the hindlimb were obtained using the attached physical markers, supplemented by functionally defined virtual markers, and detailed 3D bone surface models. A full detailed description of the methodology used to define these virtual markers and track 3D skeletal motion using a combination of XROMM data, bone surfaces and virtual markers is provided in the supplementary materials section (S1, S2).

An inverse kinematics analysis was then carried out in OpenSim [22, 32, 37] to map the kinematics from the XROMM data to the constrained biomechanical model using physical and virtual markers placed at their known locations on the model bones. Across the two kinematics trials, the ranges of motion for the skeletal kinematics with respect to hip flexion/extension (FE), abduction/adduction (ABAD) and internal/external rotation (IER) recovered in that manner were 42° (−4° to 38°), 15° (−37° to −22°), and 20° (8° to 28°), respectively. Knee and ankle FE range of motion were 40° (123° to 163°) and 74° (61° to 135°), respectively (see supplementary figure SF1). Substantial internal/external rotation range of motion at the knee and ankle was also measured, with a range of motion of 16° (5° to 21°) and 30° (−15° to 15°), respectively.

The take-off velocity of the bird was defined by measuring the velocity of its centre of mass at the instant the feet left the ground [2, 3, 6]. To that end, the centre of mass of the zebra finch was determined using the soft tissue envelope of the zebra finch and the RigidBodyParams [43] function under the assumption of a homogenous body soft tissue density [43]. The velocity of the centre of mass of the zebra finch was then measured by tracking its location throughout the take-off trials using the OpenSim 4.1 BodyKinematics Tool. Take-off velocity was determined as the velocity at the instant all toes on the hindlimb being tracked, left the perch. Take-off velocities for the two trials investigated were determined to be 1.39 and 1.08 m/s.

#### Kinetics

Previously reported ground reaction forces of the take-off leap of a zebra finch [1] were used as inputs to the current study in this study. Since full details for the collection of these data were previously reported, the experimental setup is only briefly summarized below. Vertical and horizontal ground reaction forces were recorded at 400 Hz from a force platform (Squirrel force plate, Kistler France, Les Ulis, France; resolution ±0.01N), to which a wooden perch, 1.5 cm in diameter, was attached. The ground reaction forces measured from nine trials of a zebra finch taking off from a perch [2] were used to drive the take-off simulations.

Each of the nine zebra finch kinetics trials were paired with the two sets of kinematics trials enabling analysis of 18 distinct take-off conditions. Here, the ground reaction forces were temporally aligned to the kinematic data such that at the instant at which the resultant ground force fell to zero was matched to the instant at which the toes left the ground. Ground reaction forces were assumed to act through the centre of the perch (Figure 3).

**Figure 3.**
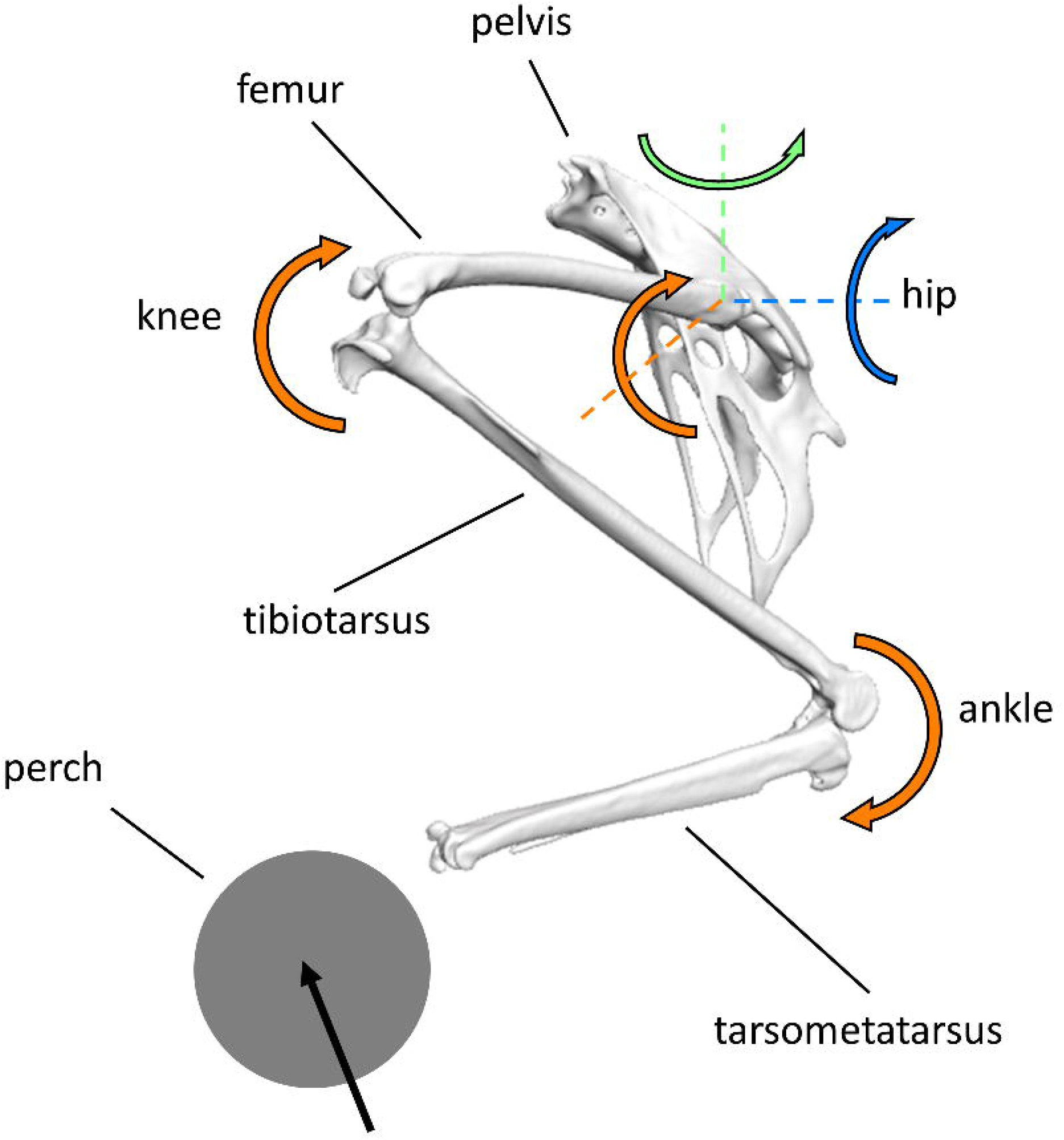
Force and moment diagram for the zebra finch hindlimb, including the pelvis, femur, tibiotarsus and tarsometatarsus in a lateral view. The musculoskeletal model described the hip, knee and ankle as three, two and two degree of freedom joints respectively. The straight black arrow on the perch represents the ground reaction force acting through the centre of the perch. Curved arrows reflect external moments acting at the joints and are colour coded such that the orange arrows identify the moment about the joint’s flexion/extension axes, blue arrows identify moments about a joints’ internal/external rotation axis, and green arrow identifies the moments about the joints’ ab/adduction axis. At the hip, the ground reaction forces consistently result in flexion, abduction and an internal rotation moments throughout the leap cycles. At the knee and ankle joints, the ground reaction forces result in extension, and flexion moments, respectively.

For use in the current study, the ground reaction force vectors were scaled in magnitude so that the impulse imparted by the leg resulted in 94% take-off velocity that was measured from the kinematics for the two trials considered here. This scaling was performed to reflect previously identified relationships for the take-off mechanics of the zebra finch [2]. Here, the method for calculating the velocity as a result of the impulse (*V_GRF_*) imparted by the hindlimb followed the approach by Provini and colleagues (2012) and is outlined below. Firstly, the acceleration resulting from the ground reaction forces, 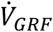, were calculated as follows:

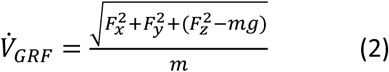

where *F_x_* is forward force, *F_y_* is lateral force, *F_z_* is vertical force, m is the mass of the bird, and g is the acceleration due to gravity. The integral of the acceleration over the duration of the take-off then provides the take-off velocity as a result of the ground reaction forces, *V_GRF_*:

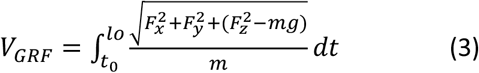

where *t*_0_ is the time when the jump starts and *l_0_* the time at lift off.

#### External joint moments

Ground reaction forces were applied to the tarsometatarsus, with the centre of pressure placed in the centre of the perch (Figure 3). Using kinematics and ground reaction forces as input to the inverse dynamics analysis, external joint moments were calculated about hip FE, ABAD, IER, knee FE and ankle FE. Joint moments were normalised by the product of bodyweight and leg length L (defined as the sum of the segment lengths of femur, tibiotarsus, tarsometatarsus, and digit III) [44], and time normalised to the duration of the jump cycle. Mean and standard deviations of the external joint moments were calculated across all 18 take-off sequences to obtain a robust estimate of the envelope of zebra finch take-off biomechanics. Here, joint moments are identified by the direction of joint movement induced by the action of the muscle group activating the respective degree of freedom.

#### Muscle moment generating capacity

The methodology for calculating muscle moment generating capacity followed the approach described by Meilak and colleagues (2021) and is outlined below. The moment generating capacity for each muscle about each joint degree of freedom (DoF) being considered was calculated by multiplying the muscle maximum isometric force (*F_max_*) by the instantaneous moment arm (*MA_t,i,j_*), evaluated at each time increment (*t*) throughout the kinematics trials. These moments were evaluated for each muscle *i (where i=1..43)* as the product of the maximum isometric muscle force (*F_max_*) and the instantaneous moment arm (*MA_t,i,j_*) determined throughout the take-off kinematics for each rotational DoF *j (where j=1..5* for hip FE, ABAD, IER, knee FE and ankle FE):

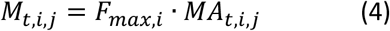

The moments of muscles acting in the same direction (i.e. their moments have the same sign) were summed for each degree of freedom to provide the total joint moment generating capacity in relation to that specific action (*M_t,j_*):

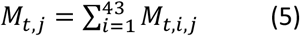

#### Comparing biomechanical conditions across the group of Passerine birds

To allow comparative analysis of conditions across the Passerine clade, the ground reaction force data available for the zebra finch was amended with similar data from further animals within the group of passerines. In a study approved by the local ethics committee of the University of Southampton (ERGO II ID 32207), a crow (*Corvus corone*) and raven (*Corvus corax*) took off 6 times each from a force platform (9260AA, Kistler) while 3D ground reaction forces (GRFs) were sampled at 10 kHz. For all species, vertical and horizontal forces were filtered using a zero-phase low-pass (40Hz) custom filter in Matlab. The peak ground reaction forces of the zebra finch, starling [3], crow and raven were all normalised by bodyweight and, together with key measures of hind limb geometry (Table 2) to compare take-off conditions within the group of passerines.

**Table 2.** Key morphometric parameters of zebra finch (Taeniopygia guttata), starling (Sturnis vulgaris), crow (Corvus corone), and raven (Corvus corax).

| species | mass<br>[g] | femur length<br>($L_{fem}$) [cm] | tibiotarsus<br>length ($L_{tib}$)<br>[cm] | tarsometatarsus<br>length ($L_{tars}$) [cm] | digit III<br>length<br>[cm] | hindlimb<br>length<br>[cm] | hindlimb<br>index<br>$((L_{tars} + L_{tib}) / L_{fem})$ |
| --- | --- | --- | --- | --- | --- | --- | --- |
| zebra<br>finch | 15.4 | 1.40 | 2.24 | 1.46 | 1.10 | 6.2 | 2.64 |
| starling<br>[3] | 77.3 | 2.53 | 4.37 | 2.82 |  |  | 2.84 |
| crow | 440 | 5.28 | 8.74 | 5.77 |  |  | 2.75 |
| raven | 1100 | 6.92 | 11.42 | 6.80 |  |  | 2.63 |

## Results

The duration of the take-off leap ranged between 62-65ms (Figure 4). Mean resultant peak ground reaction forces per leg were 1.81±0.21 BW (range: 1.42 to 2.15 BW, c.f. supplementary figure SF1) and occurred at about 62 % leap cycle time. Across the two kinematics trials, the largest ranges of motions were 74°, 42° and 40° about the ankle, hip, and knee FE axes, respectively. The ranges of motion about the joints’ secondary degrees of freedom were similar, with hip IER and ABAD RoMs of 20° and 15°, respectively and knee and ankle IER RoMs of 16° and 30°, respectively (see supplementary figure SF2).

**Figure 4.**
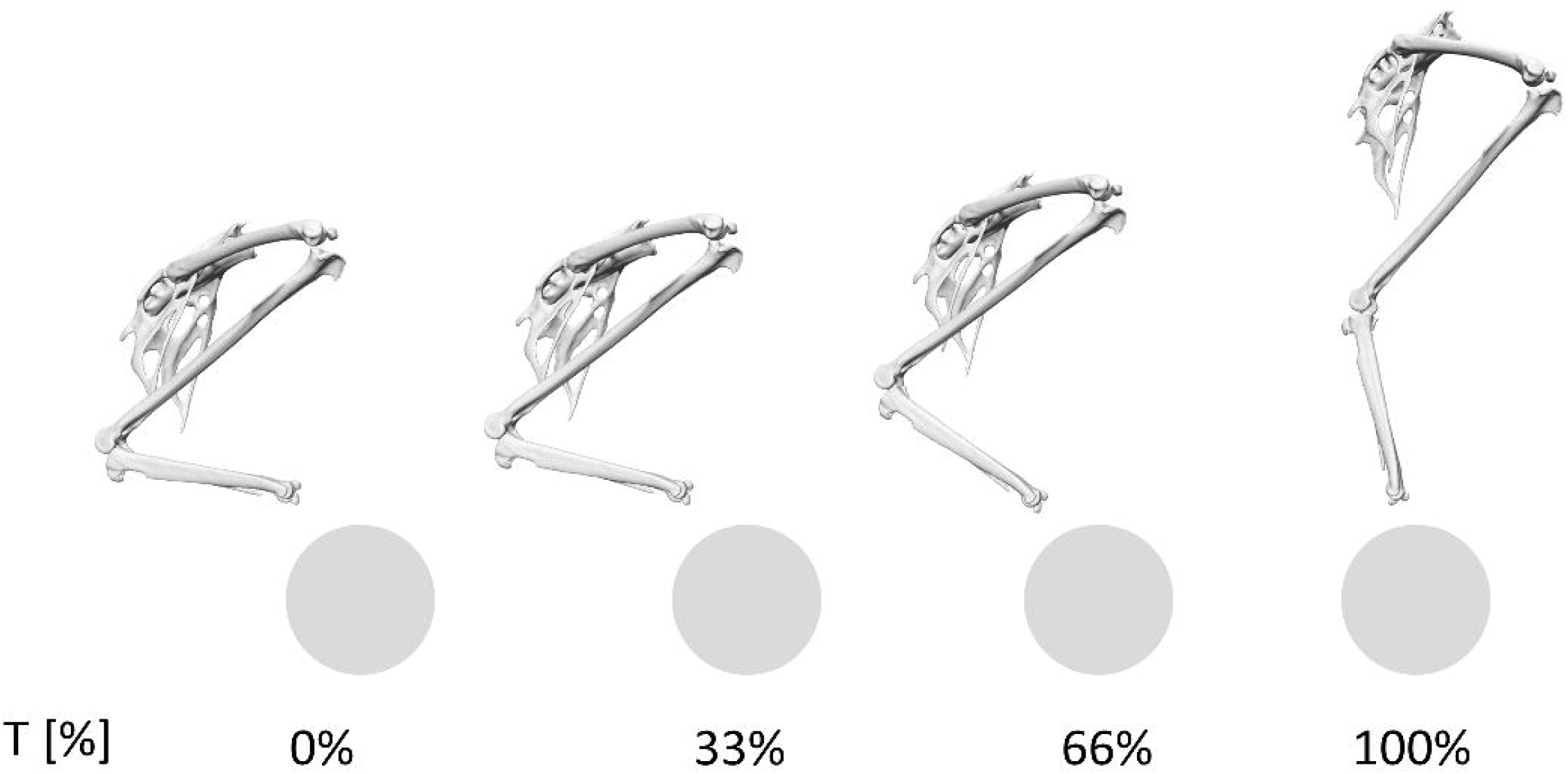
Lateral view of right zebra finch hindlimb throughout the normalised take-off leap cycle. The duration of the take-off jump duration ranged from 62 to 67 ms while all of the ankle, knee, and hip joint undergo substantial extension.

The largest external moments observed were the moments around the flexion/extension axes. Here, the mean peak external joint flexion moments at the hip and ankle were of similar magnitude (0.66±0.04 and 0.68±0.05 BWL, respectively) while the mean knee extension moment was only about half that value (0.38±0.05 BWL, Figure 5, Table 3). The smallest peak moments were computed about hip IER and ABAD with mean peak moments of 0.27±0.07 and 0.20±0.07 BWL, respectively (Figure 5, Table 3). At the instances that these peak moments occurred, the muscles had the ability to generate 120%, 282% and 61% of the mean peak hip FE, IER and ABAD joint moments, respectively (Figure 5 Table 3). The muscles were able to generate 177% and 176% of the mean peak joint moments about the FE axes of the ankle and knee, respectively (Figure 5 B and C, Table 3).

**Table 3.**
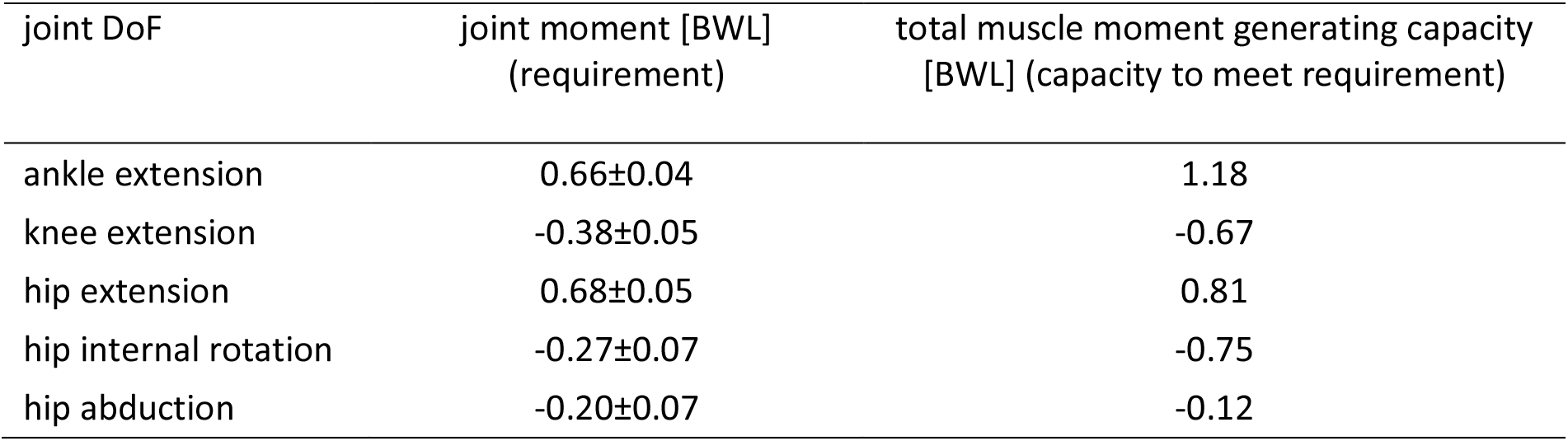
External joint moments and total muscle moment generating capacities of the zebra finch at the instances at which the peak external joint moments occur. To actively power the take-off leap by muscle action, the combined(total) moment generating capacity of all muscles must at least reach if not exceed the level of the external joint moments. Positive moments act in extension, external rotation, and adduction, while negative moments act in flexion, internal rotation, and abduction.

**Figure 5.**
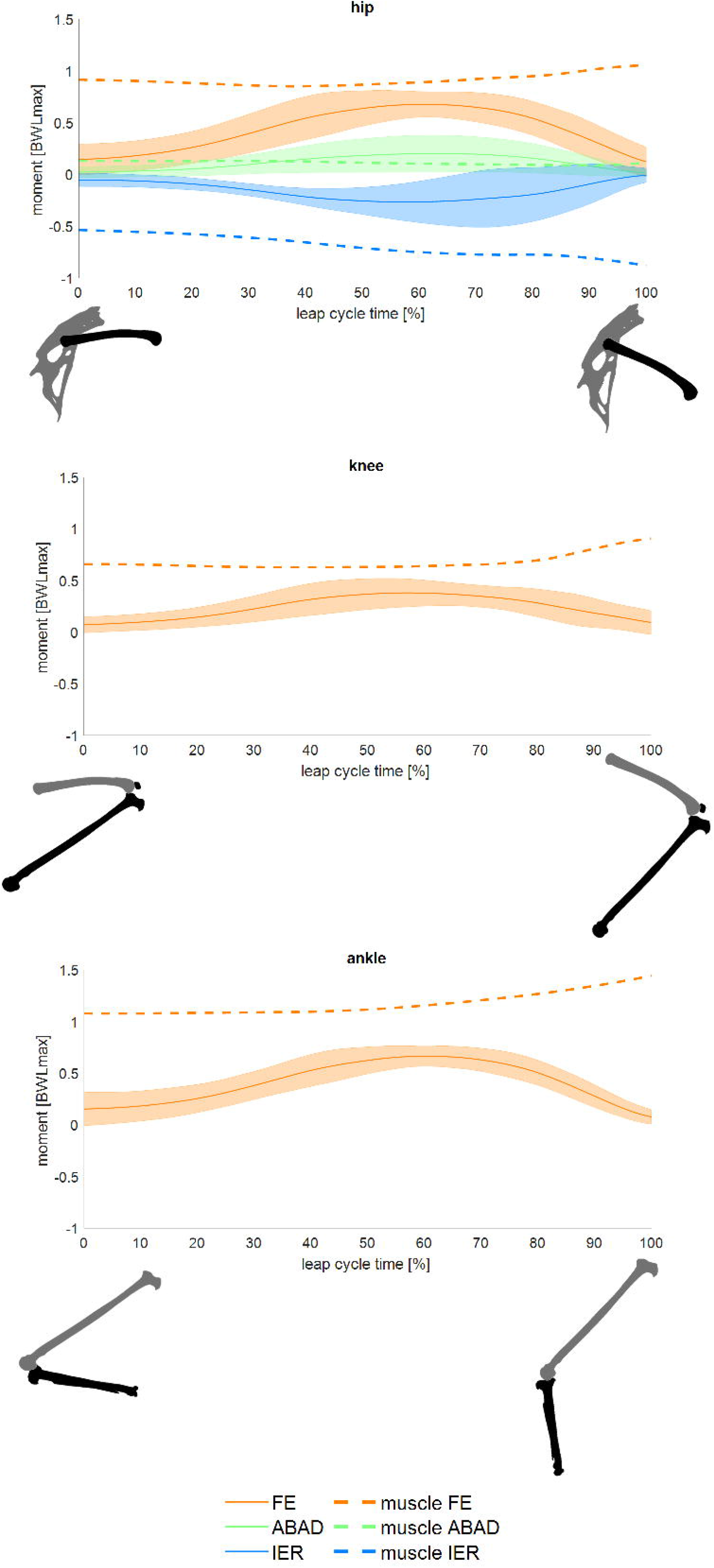
External joint moments at the hip, knee and ankle together with the total moment generating capacity of the muscles, plotted over normalised leap cycle time. Solid lines depict the mean external joint moments (requirements), bands depict ±2.5 standard deviations, while dashed lines depict the total moment generating capacity of the muscles (capacity). Colours are used to differentiate the axis around which the moments act, with orange representing flexion/extension, green ab/adduction, and blue internal/external rotation. Positive moments represent (internal/external) moments that act to extend, adduct, and internally rotate, respectively.

Comparing conditions across the group of passerines, the zebra finch as bird with the smallest mass by far (15.4 g) had the smallest normalised peak ground reaction forces per leg of 1.94±0.25 BW, while starling (77.3 g), crow (440 g), and raven (1.1 kg) exerted very similar peak GRFs with values of 2.15±0.14 BW [3], 2.20±0.29 BW, and 2.25±0.14 BW, respectively (see supplementary figure SF3).

## Discussion

This is the first detailed investigation into the hindlimb mechanics of a flying bird as it takes to the air with a take-off leap using modern biomechanical analyses. The model presented here simulated 18 distinct take-off trials of a zebra finch’s take-off leap from a perch covering a range of take-off conditions in a passerine bird, the group that makes up approximately 60% of all extant bird species [7–9]. In doing so our study revealed a consistent pattern of internal mechanics across the trials that was characterised by the largest peak external moments occurring around the FE axis of the hip and ankle (requirements, 0.68 and 0.66 BWL, respectively), while considerably smaller external moments were observed about the FE axis of the knee, amounting to only 56% of the respective peak moments at the hip. On the other hand, peak hip IER and ABAD moments were 40% and 29% of peak hip FE moments respectively. Together with previous findings regarding the substantial extent of joint range of motion in IER exercised by birds [1, 27] and the substantial moment generating capacity of the hip joint muscles around that axis [10], these novel data on the internal hind limb mechanics during the take-off leap strongly support the notion that the execution and control of hindlimb motion in birds is not limited to a single (sagittal) plane but is essentially 3D in nature [16].

Across the hip, knee and ankle FE degrees of freedom, the muscles had the ability to generate 120%, 176%, and 177% of the mean peak external joint moments, respectively. In relative terms therefore, muscles spanning the hip were closest to reaching the capacity limit whilst muscles spanning the knee and ankle joints had the largest reserves. Although for the most typical (mean) of the conditions studied here the capacity of the muscles to generate moments was always larger than the requirement (Figure 5), the closing gap between the upper limit of the 2.5 SD range of the external hip flexion moments and available hip muscle extension capability suggests that more powerful take-off leaps than those observed here would likely require further activation of the more distal, knee and ankle joint spanning muscles for which the requirements remained more comfortably within their capability [23]. An explanation as to why capacity to balance the moments at the more distal joints retains a larger reserve may be found in the design of the avian hindlimb and specifically the biarticular muscles [45–47]. Such biarticular muscles include in particular the m. caudofemoralis pars caudalis (MFCLP), m. flexor cruris medialis (MFCM) and m. iliofibularis (MIF) which span the caudal side of the hip and knee (Figure 1, Table 1). MFCLP and MFCM form a major part of the hip extensor moment generating capacity, together generating 35% of the total hip extensor moments whereas the MIF has a greater role as a hip abductor, generating 20% of the muscle abduction generating capacity [10]. Whilst during the take take-off leap of the zebra finch the hip, knee and ankle are all extending (Figure SF1), the muscles need to generate a net knee flexion moment, suggesting that knee flexors may be activated. Biarticular muscles, which if activated generate an extension moment at the hip but a flexion moment at the knee such as the MFCLP and MFCM, would thus appear to be prime candidates for meeting the demands during the take-off leap. Moreover, co-activation of knee flexors and extensor muscles (muscle co-contraction) though energetically less optimal, may help to increase compressive forces across the knee joint and help to lock or at least minimise the extent of internal/external rotation [27, 28, 48]. Further analysis to examine muscle activation patterns, though beyond the scope of the current work, could corroborate whether activation of these biarticular muscles to extend the hip and generate the extension moment applied to the knee suggested by the analyses here does indeed occur and help to further elucidate how avian hindlimb muscles are orchestrated during the take-off leap.

Here, it is also informative to consider absolute moments where indeed the ankle extensors had the largest joint moment generating capacity, capable of generating peak total extension moments of up to 1.18 BWL at the ankle, followed by the hip extensors, with a peak total capacity of 0.81 BWL. The passerine bird ankle muscles’ capacity to produce the largest moments indicates the importance of the ankle joint throughout the take-off, a finding in line with previous research on the ground dwelling elegant-crested tinamou *Eudromia elegans* [23] where the take-off heavily depended on the parameters and activation of the ankle extensors. Though to the authors knowledge no similar studies reporting internal mechanics during a take-off leap of a flying bird are available for comparison, similar analyses have been performed in ground-dwelling birds such as the emu and the ostrich. Here, data on the running of the ostrich obtained using similar analysis methods revealed that the peak normalised FE moments across all of the hip, knee, and ankle joint of the ostrich were considerably smaller than those reported during the take-off leap of the zebra finch reported here. The largest differences in normalised moment magnitudes were observed for the ankle and hip joints where the peak moments during running in the ostrich amounted to only 24% (0.16 BWL) and 25% (0.17 BWL) of the values for the zebra finch leap. The external moment at the knee during running in the ostrich was the largest of all 3 hindlimb joints with 0.20 BWL yet amounted to only about 53% of the value the current study calculated for the take-off leap of the zebra finch. In the comparison of the absolute moment magnitudes it is important to consider that speed at which the ostriches were running was rather slow (3.24 m/s) compared to the maximum speed ostriches can achieve (about 13.9 m/s) and higher speeds will be associated with higher external forces and moments. However, not only absolute magnitudes but also the ratios of their magnitudes at the hip, knee, and ankle differed between the zebra finch take-off leap and ostrich running. While the largest external moment during running of the ostrich was computed at the knee, suggesting that that ostrich running is knee driven, the external knee moment during the take-off leap of the zebra finch was the smallest of all the hind limb joints. For the zebra finch leap, peak moments were observed at the hip and ankle suggesting that this motion is hip and ankle driven instead and signifying that an interesting avenue for future work would be to investigate whether the different motor behaviours are indeed associated with different muscle coordination patterns and that care must be taken when speculating about design principles and interpreting optimality of the musculoskeletal system based on a limited repertoire of motor behaviours.

Previous studies have demonstrated that avian pelvic muscles have a considerable capacity to produce IER moments at the hip [10, 16, 49]. The current study revealed not only that substantial external moments about the IER axis occur during the take-off leap of the zebra finch, approaching 40% of the hip extension moment, but also demonstrated that the hip muscles had the greatest relative capacity to actively balance these moments, with muscles capable of generating up to 280% of the mean peak external IER moments. Together these data provide further evidence that IER rotation is actively controlled during routine, straight line take-off leap of passerine birds. The ample capacity of the muscles to enable IER during such jumps further allows for take-off leaps to occur with substantially more axial turning/rotation while the bird remains on the ground, offering the bird a wider repertoire of take-off and escape behaviours. On the other hand, foraging behaviours have been shown to be linked with substantial extents of hindlimb IER [27] and it may well be that substantial IER capability of the hip muscles are particularly crucial in supporting those behaviours (particularly for single limb support) in addition to allowing variation in the take-off leap.

In contrast to the well powered hip IER DoF during the take-off leap, the pelvic muscles of the zebra finch were only able to balance 61% of the peak mean external adduction moments. Analysis of the relationship of the relative moment generating capacity of the avian hip muscles do indeed demonstrate that the smallest capacity to generate a moment exists with respect to the ABAD axis. However, even though the avian hip has a relatively limited capacity to actively produce hip ABAD moments, birds can rely on a passive mechanism using the antitrochanter and associated ligamentous structures to balance external abduction moments [49]. The utilisation of the antitrochanter is also a feature in the hip mechanics of the ostrich: though substantial external abduction moments are generated during walking and running in these flightless birds, the design of the hip allows to stabilise the joint passively such that ostriches neither require nor possess muscles to do so actively [49, 50]. The antitrochanter is indeed a feature shared across all extant birds that was not present in some of the very earliest birds such as *Archaeopteryx*, who likely relied on powerful hip adductors to generate the required adduction moments [16].

Maintaining substantial lever arms of the muscles throughout the functional range of motion of a joint is a prerequisite to maintain high levels of muscle capacity to generate moments. Passive structures can play a key role to help maintaining muscle lever arms include sesamoid bones such as the patella which is key to enable joint function in flexion at the knees in human and avian bipeds [33, 46] The hypotarsus is another anatomical feature of the avian hindlimb that helps to guide tendons around the ankle joint and to and maintain their lever arms [51]. The posterior side of the ankle further includes the cartilagio tibialis which constrains the muscle line of action to act further away from the joint axis of rotation thus helping to maintain its moment [29]. The model presented here therefore modelled the patellofemoral joint in an approach informed by the birds’ bone surface anatomy and kinematic model data from the literature [38] and in incorporated these passive structures of the ankle with wrapping objects and via points informed by CT scans.

The relative peak ground reaction forces during the take-off leap of all passerines considered here were rather similar in magnitude, despite the large range in body mass. During the take-off leap the zebra finch (15.4 g), starling (77.3 g), crow (440 g) and raven (1.1 kg) generated peak ground reaction forces per leg of 1.94±0.25 BW, 2.15±0.14 BW [3], 2.2±0.29 BW, and 2.25±0.14 BW, respectively. Similarities do not stop with the ground reaction forces but are also apparent in the morphology of their hindlimbs. The passerines considered here, the zebra finch, starling, crow, and raven, possess very similar hindlimb indices, a measure of relative segment lengths of the hindlimb ((tarsometatarsus length + tibiotarsus length) / femur length [52]) with values of 2.64, 2.84, 2.75, and 2.63, respectively [53] (Table 2). The similarities in normalised leg segment lengths and ground reaction forces spanning a range of passerines, which differ in mass by approximately two orders of magnitude, support the hypothesis that passerines share a similar take-off behaviour, as reported here. Forward dynamics simulations of the tinamou (0.55 kg) leaping predict similar but somewhat higher peak ground reaction forces of 2.62 BW per leg during the jump [23]. On the other hand, predicted joint kinematics profiles for the take-off leap of the tinamou demonstrated ranges of motion at the hip knee and ankle of 65°, 91° and 109° respectively, consistently greater than the range of motion measured in the zebra finch (42°, 40° and 74° respectively), while the hindlimb index of the tinamou *Eudromia elegans* (belonging to the group Tinamiformes) was 2.21, substantially smaller than that of the passerines considered here. The relative difference in relative leg morphology between the passerines and tinamou could be one of the determining factors resulting in the variability in take-off mechanics between the clades of birds.

The kinematics used to drive the take-off simulation were informed by previously obtained XROMM data of the zebra finch take-off leaps [1]. In this study, the use of the tantalum bead markers, detailed bone surface geometry from high resolution μCT, and anatomical-functional relationships [31, 54] were all used to increase repeatability in tracking 3D skeletal kinematics and reduce the influence of the user during scientific rotoscoping [42]. Across the two kinematics trials, the difference between the originally published orientation angles of the femur, tibiotarsus and tarsometatarsus was smallest about FE and ABAD axes of the bones, with mean differences ranging from 0.6±0.5° to 3.3±2.7°. The degree of freedom most difficult to determine during scientific rotoscoping was IER due to the cylindrical shape of the long bones. Predictably, the largest difference was observed when comparing the bone IER orientations, with mean differences in IER orientations across the three long bones ranging from 7.3±4.8° to 11.5±6.3°. Differences in bone locations between the two methods were minimal, mean differences across all three long bones ranged between 185 – 333 microns.

Due to the limited number of specimens in which sufficient bones had had a minimum of three tantalum markers attached, only two sets of kinematics trials were used in this study. However, by pairing each set of kinematic trials with nine sets of kinetics trials, we maximised the variability in take-off conditions studied here. Future studies, using XROMM to capture detailed kinematics of passerines should ensure that at least one of the long bones includes at least 3 markers to reduce the reliance on the user during scientific rotoscoping. The simulations here included IER motion of the knee and ankle joints, however the active muscle actuation of these degrees of freedom was not considered in line with previous studies and under the assumption that typically small moments are balanced by passive structures such as ligaments [37, 55]. Previous studies reported the take-off velocity of the zebra finch to be around 1.7m/s [2, 6] which is faster than the take-off velocity, measured here from the XROMM data which ranged between 1.08-1.39 m/s. Previously published studies measuring the kinematics of animal subjects using implanted markers have reported the markers causing a limp [14]. The comparatively slower take-off velocities reported here could thus be attributed to a response to the implantation of the tantalum bead markers on the hindlimb bones. However, varying levels of motivation between experimental conditions may also explain the observed difference and the similar protocols using tantalum bead markers have been used to study a range of motor behaviours in birds [46, 56, 57]. This study did not include the mechanics of phalanges in the analysis, as the take-off trajectories of the hindlimb are defined primarily by the motion at the more proximal hindlimb joints where also more substantial joint moments are generated. Future studies which may consider hind limb mechanics during landing, when the detailed mechanics of foot are likely to play a more important role, should aim to capture the detailed kinematics of the phalanges.

This study considered the maximal moment generating capacity of the muscles, taking into account the muscle maximum isometric force and instantaneous moment arm throughout the take-off leap and contrasted these to the external moments applied to the joints of the hind limb. In this way general patterns of mechanical requirement and hindlimb muscles capability to meet the requirements of a take-off leap were analysed. Although the sensitivity of muscle moment generating capacity with respect to uncertainty in the definition of muscle geometry was not directly investigated in this study, a thorough sensitivity analysis was performed for pelvic muscles in the magpie hindlimb model that was the basis for the current study [10], which, given the similarity in overall hindlimb design and specific hindlimb bone morphology, can reasonably be expected to remains valid for the musculature of the zebra finch model here. Though the determination of the detailed muscle activation patterns to balance the external moments was not within the scope of the current study, further analyses of the biomechanical model (such as static optimisation [15, 32, 37]) would help to further elucidate the detailed activation patterns of individual muscles as well as providing estimates for the likely bon-on-one joint contact forces being transferred at le large joints of the avian hind limb during a take-off leap.

This study is the first to establish the biomechanical requirements of the hindlimb of a flying bird as it takes to the air. We present biomechanical conditions that hindlimbs of passerines, a clade of birds that include over half of all avian species, experience during take-off. The zebra finch take-off leap is primarily hip and ankle driven, which is in direct contrast to ostrich running mechanics which indicates avian running is a primarily knee driven activity. The ability of the zebra finch muscles to produce over double the mechanical requirements at the ankle and knee axes and about 20% more than the requirements about the hip FE axis along with the suspected use of biarticular muscles and passive structures is consistent with the hypothesis that the take-off leap as reported here is an optimized way for the zebra finch to take to the air. Striking similarities in ground reaction forces and relative leg morphology of multiple passerines suggest that the take-off behaviour described here could be shared across all passerines, despite differences in mass by two orders of magnitude.

## Acknowledgements

We would like to acknowledge the skill, hard work, and patience of the bird handlers Lloyd and Rose Buck who helped with the collection of the ground reaction force data of the crow and raven. This work was supported by the Natural Environmental Research Council [grant number NE/L002531/1] and grants from the UMR 7179, l Action Transversale du Muséum National d Histoire Naturelle formes possibles, formes réalisées and from Ecole Doctorale Frontières du Vivant and Bettencourt-Schueller Foundation fellowships as well as the National Science Foundation [grant nos IOS-0923606 and IOS-0919799]. Travels were paid by the UMR 7179.

