## Supplementary information for "On the hindlimb biomechanics of the avian take-off leap"

### S1 Optimised tracking of skeletal kinematics

In order to reliably track the 3D skeletal kinematics for use in the musculoskeletal analyses, bone surfaces were mapped from the CT space to the XROMM space using a combination of physical and virtual markers while additionally minimising penetration of the hindlimb bone surfaces as detailed below.

#### S1A Functional – anatomical definition of virtual markers

The 3D positions of the physical markers which were tracked throughout the leaping motion using XrayProject 2.2.4 in MATLAB [1, 2] formed the basis for mapping the bone surfaces from the CT system to the XROMM space using Ordinary Procrustes Analyses (OPA). To enable mapping of the surfaces using OPA, knowledge of the location of at least 3 markers was required for each tracked bone. Since the tarsometatarsus was the only bone to be tracked on which 3 physical markers were placed in the experiment, additional virtual markers were defined at the hip, knee and ankle joint centres. The definition of these virtual markers commenced by calculating functional axes of rotation [3-6] of the ankle, knee and hip using the µCT data and the bone surfaces derived from that data. After confirming minimal morphological difference between bone surfaces of the left and right hindlimbs, the surface of the pelvis was mirrored at its mid-sagittal plane [7] and registered to its original shape using rigid ICP registration [8]. The right femur was then mirrored similarly and mapped with the registration matrix previously established for the pelvis. The left femur was then registered to the mirrored and mapped right femur, eventually providing two different poses for a left femur with respect to the left acetabulum. Using these two joint poses, the functional axis of rotation of the left hip was calculated from the vertex positions of the respective bone surfaces (figure S4 A) [5]. Spheres were fitted to the articulating surfaces of the femoral heads (in two poses) and acetabulum (Mesh2Surface for Rhino v6.1.5 in Rhino v7) [9]. The centres of these spheres were projected on to the functional axis of rotation of the hip and their mean position was taken as the functional-anatomical hip centre of rotation (figure S4 A). This hip centre could be expressed with respect to both the pelvis and the femur, providing a virtual marker for both bones and an objective means to link them via a ball and socket joint. A similar approach was used for functionally defining the knee and ankle joint centres. The right femur was mirrored and registered to the left, again using rigid ICP registration [8]. Using the transformation matrix resulting from the aforementioned registration, the right tibiotarsus was mapped to the left hind limb before the left tibiotarsus was ICP-registered to that pose, resulting in two poses of the left tibiotarsus with respect to the left femur (figure S4 B). Using these two joint poses, the functional axis of rotation of the knee was calculated [5]. The midpoint of the intersection of this functional axis with the surface of the femur defined the functional-anatomical knee joint centre (Fig. S4 B). In a similar manner, the mirrored right tibiotarsus was registered to the left, and that registration was used to map the mirrored right tarsometatarsus to the left before ICP-registering the left tarsometatarsus to that pose (Fig. S4 C). Using the two resulting joint poses, the functional axis of rotation of the ankle was found [5]. The midpoint of the intersection of the resulting axis of rotation with the tibiotarsus provide a further virtual marker at the functional-anatomical ankle centre.

### S1B Collision detection supported reconstruction of skeletal kinematics

After augmenting the physical markers with the virtual markers located at the hip, knee and ankle joint centres, reconstruction of skeletal kinematics proceeded with mapping bone surfaces from the space of the CT system to the XROMM system by OPA between markers associated to each bone. Given that 3 physical markers were attached to the tarsometatarsus only, the surface mapping process started from that bone. OPA computed between the 3 physical markers attached to the bone in the CT and XROMM systems was used to map the tarsometatarsus surface and also the ankle joint centre (virtual marker) from the CT to the XROOM. Further OPA calculated from the virtual ankle joint centre and the two physical markers of the tibiotarsus and their respective µCT location was then used to map the tibiotarsus surface and the knee joint centre (virtual marker) from the CT to the XROMM system.

As only a single physical marker was attached to the femur, at this stage of the process the position of only two femoral markers (1 virtual, 1 physical) were known in the XROMM space. To determine the position of a third marker that maximised the use of the XROMM marker data in a manner consistent with the kinematic model of a ball and socket joint for the hip, a virtual hip joint centre marker was derived as follows. Using the previously determined functional anatomical hip centre it was possible to construct two spheres, centred at the location of the respective physical markers of the femur and pelvis, respectively, with radii corresponding to the previously determined distance between these physical markers and the functional-anatomical hip centre. By determining the intersection between these spheres, a circle could be determined on which the hip centre was known to lie. The exact location of hip centre on the circumference of the circle was determined by preventing any collision between the surfaces of the tibiotarsus and the distal femur while also minimising the extent of collision between the surfaces of the femur and the pelvis (acetabulum and anti-trochanter). The implementation of this process in Matlab (2019b, The Mathworks, Nantucket, USA) sampled the circumference of the circle on which the hip centre was known to lie in 1 degree intervals to find the pose of the femur for which there was no intersection with the surface of the tibiotarsus and minimal intersection with the surface of the acetabulum. At the final stage of the process, the maximum intersection volume remained at very small values at below 0.37% of the volume of the femur.

### S2 Definition of anatomical bone and joint coordinate systems

Anatomical coordinate systems for the bones and joint coordinate systems (ACS and JCS, respectively [10]) were defined for the long bones where ACSs were defined at the proximal end of long bones whilst the JCSs were defined at the distal portion of the bone. Orientation of the coordinate systems followed conventions described in the literature [2, 10, 11] (Fig. S5). The origin of the pelvic ACS was defined at the midpoint between the two spheres fitted to isolated regions of the acetabular joint surfaces using the methods of least squares (lssphere.m v1.0, Matlab (2018a, The Mathworks, Nantucket, USA)). A midsagittal plane was defined using an approach where the mirrored pelvis surface was registered to its original shape [7]. The Z axis of the pelvis ACS was the normal of the midsagittal plane pointing from right to left and the Y axis (positive axis direction pointing dorsally) was derived from a projection of the 1$^{st}$ principal axis of the pelvis [12] on the midsagittal plane. The pelvic ACS X axis was determined from the cross product of the Y and Z axes, with the positive axis pointing cranially. The axes of the JCS of the left hip were aligned with the axes of the pelvic ACS while its origin was located at the respective functional-anatomic hip centre of rotation determined previously.

The X axis direction of the femoral ACS (positive direction pointing proximally) was defined by determining the centroid line of the femoral shaft to which a straight line was fitted using the method of least squares (lls3dline.m v1.0). Its Z axis (positive direction pointing laterally) was defined between the femoral head centre and its projection on to the fitted shaft axis. The Y axis of the femoral ACS (positive direction pointing anteriorly) was determined from the cross product of the X and Z axes. The functional-anatomical knee centre of rotation, determined previously, was defined as the origin of the femoral JCS. The functional knee axis of rotation defined the Z axis of the JCS (positive axis direction pointing medially). The Y axis of the femoral JCS (positive axis direction pointing anteriorly) was determined from the cross-product of the Z axis of the JCS with the X axis of the ACS. The X axis of the femoral JCS (positive axis pointing proximally) was determined from the cross-product of its Z and Y axes.

The processes for defining the local coordinate systems of the tibiotarsus and tarsometatarsus were similar: the ACS X axis direction (positive axis pointing proximally) was defined by least squares fitting a line to a centroid axis of the bone shaft. The Z axis of the JCS (positive axis pointing laterally) of the tibiotarsus was defined using the ankle axis of rotation whereas for the tarsometatarsus the Z axis was defined between the points of intersection of cylinders fitted to the distal medial and lateral condyles with the bone surface (lscylinder.m v1.0). The Y axes (positive axis pointing anteriorly) of both the ACS and JCS were defined by calculating the cross product between the JCS Z and ACS X axes. JCS X axes (positive axis pointing proximally) were defined by calculating the cross product between the JCS Y and Z axes. ACS Z axis (positive axis pointing laterally) was defined by calculating the cross product between the ACS X and Y axes. Tibiotarsal and tarsometatarsal ACS origins were defined at the intersection of the shaft axes with the proximal surface of the respective bone. The functional-anatomical ankle joint centre of rotation was taken as the origin of the JCS of the tibiotarsus.

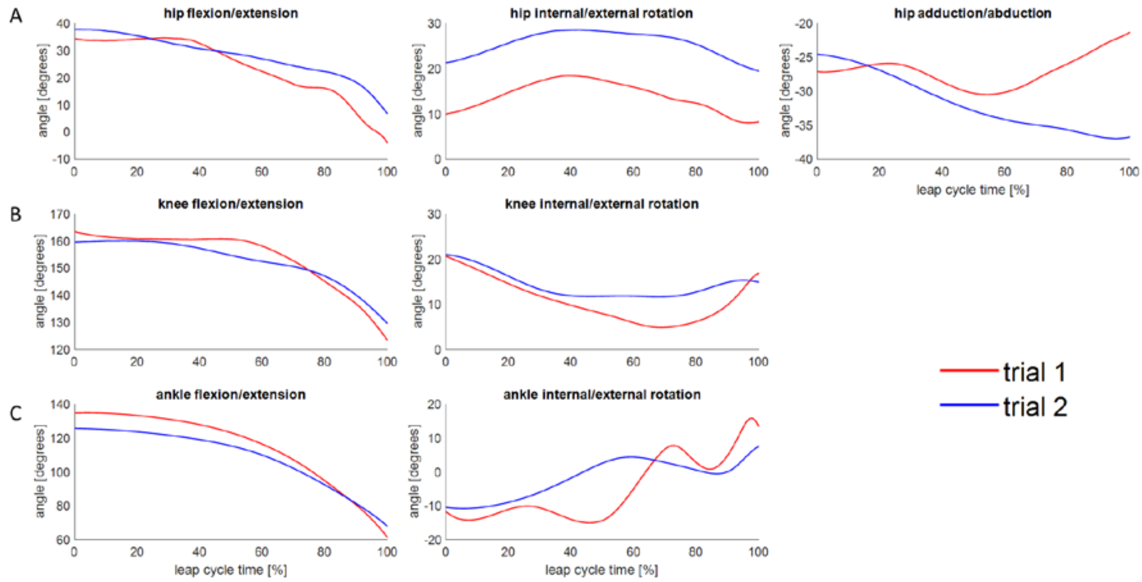


Figure SF1. Joint angles at the A) hip, B) knee and C) ankle for 2 take-off trials derived from XRoMM data. Hip joint flexion/extension (FE), internal/external rotation (IER), abduction/adduction (ABAD), knee, and ankle FE and IER are shown here where positive values for FE, ABAD and IER are flexion, adduction, and internal rotation, respectively. Line colours represent the specific trial from which the data was derived (red: trial 1; blue trial 2).

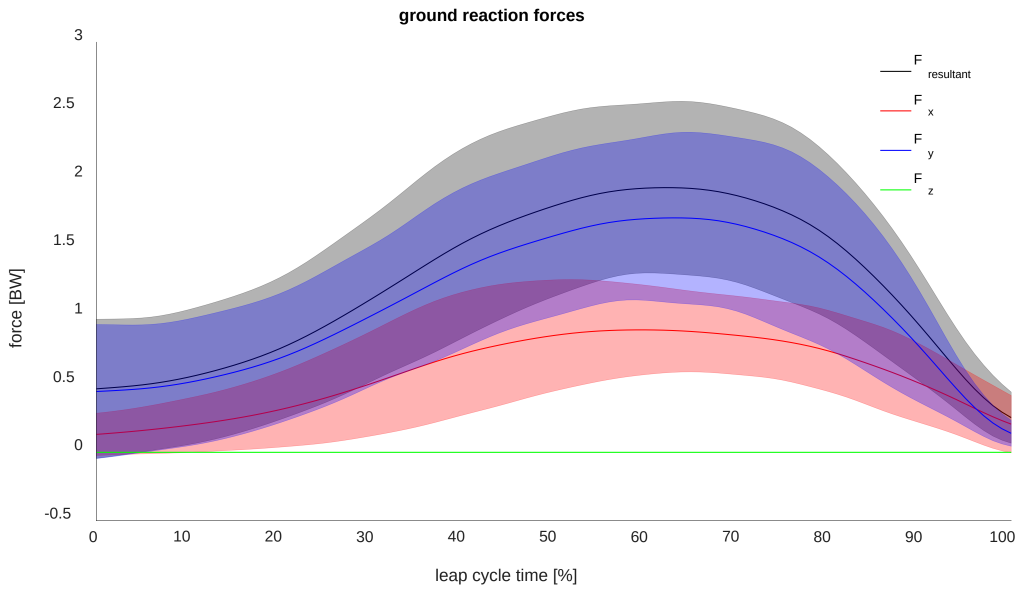


Figure SF2. Resultant and individual x, y, z components of the ground reaction force during a full take-off cycle. Solid lines represents mean values while shaded bands surrounding the mean represent ±2.5 standard deviations (SDs). Force components Fx, Fy, and Fz are horizontal (caudal-cranial), vertical (ventral-dorsal), and sideways (medial-lateral) directions, respectively.

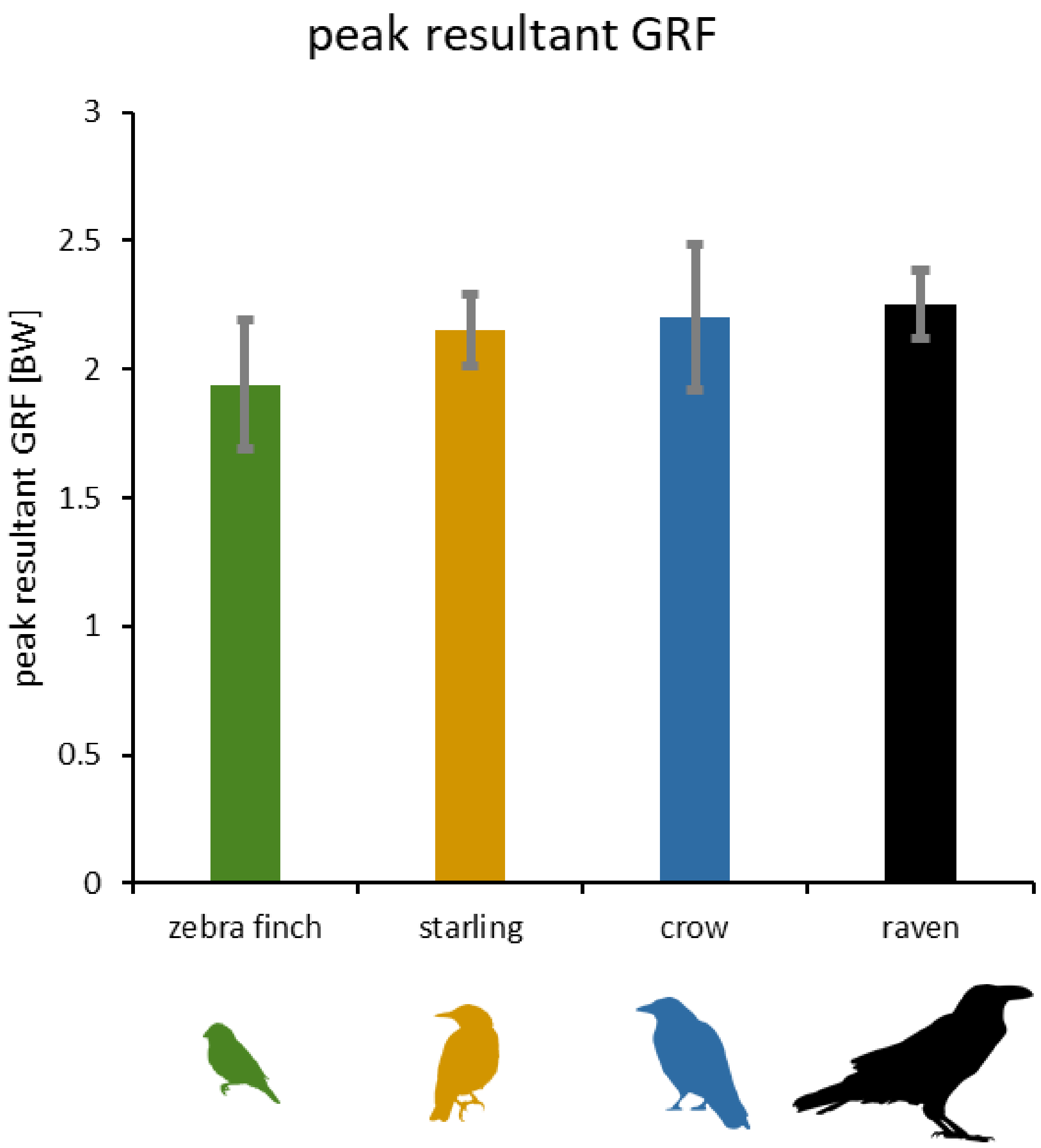


Figure SF3 Mean ± standard deviation of the peak resultant ground reaction forces (GRFs) acting on one leg of the zebra finch, starling [13], crow, and raven during their respective take-off leaps. Silhouettes from phlyopic.org.

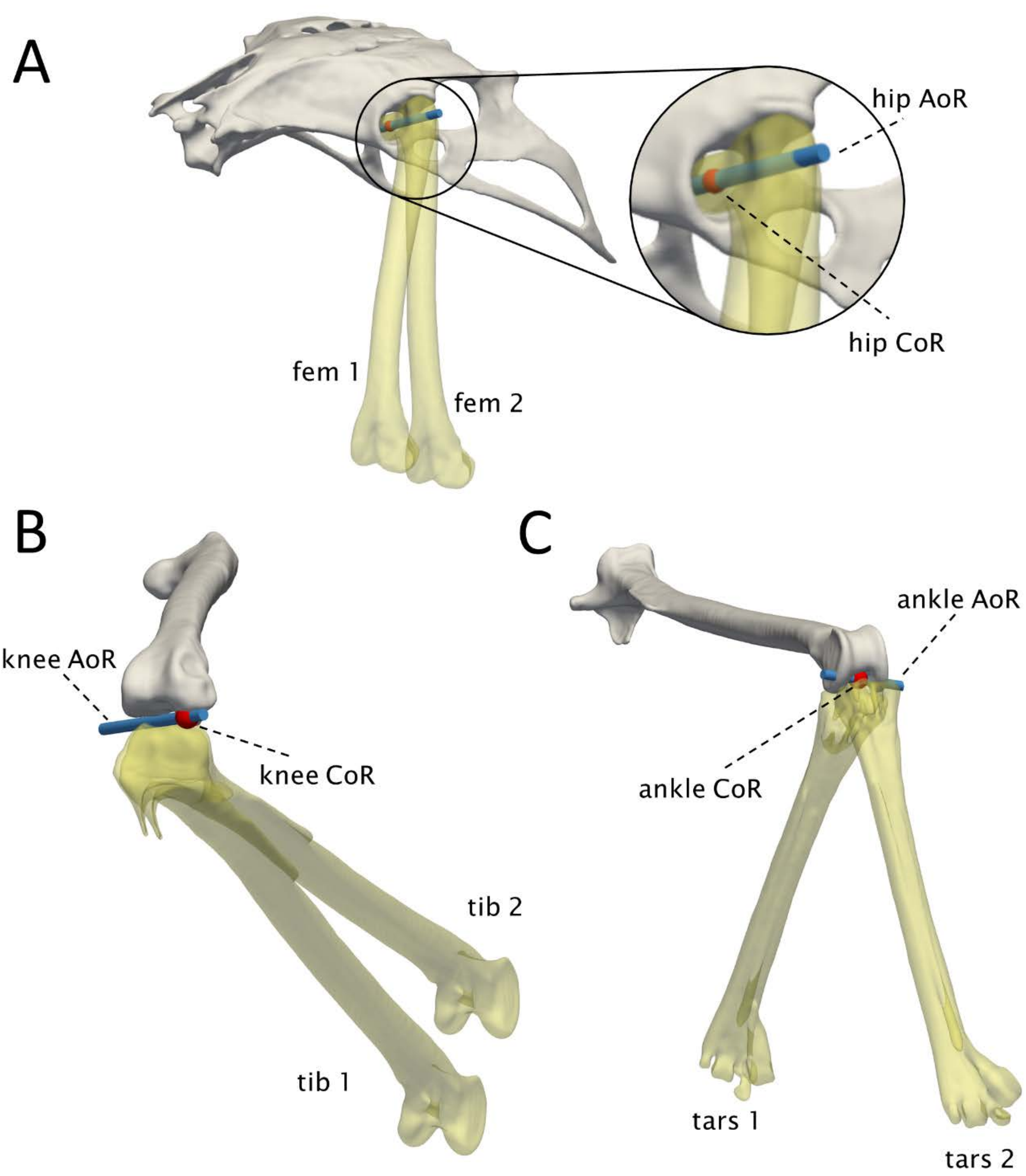


Figure SF4 Functional anatomical joint centres for the hip, knee and ankle joint were derived from a CT scan of the zebra finch hindlimbs. For each joint, the difference in pose between the left and right hindlimbs was used to estimate a functional joint axis of rotation (AoR) while joint centres were derived from the intersection of the AoR with the surface of the proximal bone. Blue cylinders represent the functionally determined joint axes of rotation (AoR) and red spheres represent the functional-anatomical joint centres of rotation (CoR). A pose of the mirrored right femur (fem 2) registered to the left side, together with the original pose of the left femur (fem 1). B mirrored right tibiotarsus (tib 2) registered to the left, together with the original pose of the left tibiotarsus (tib 1). C pose of the right tarsometatarsus (tar 2) registered to the left, together with original pose of the left tarsometatarsus (tar 1).

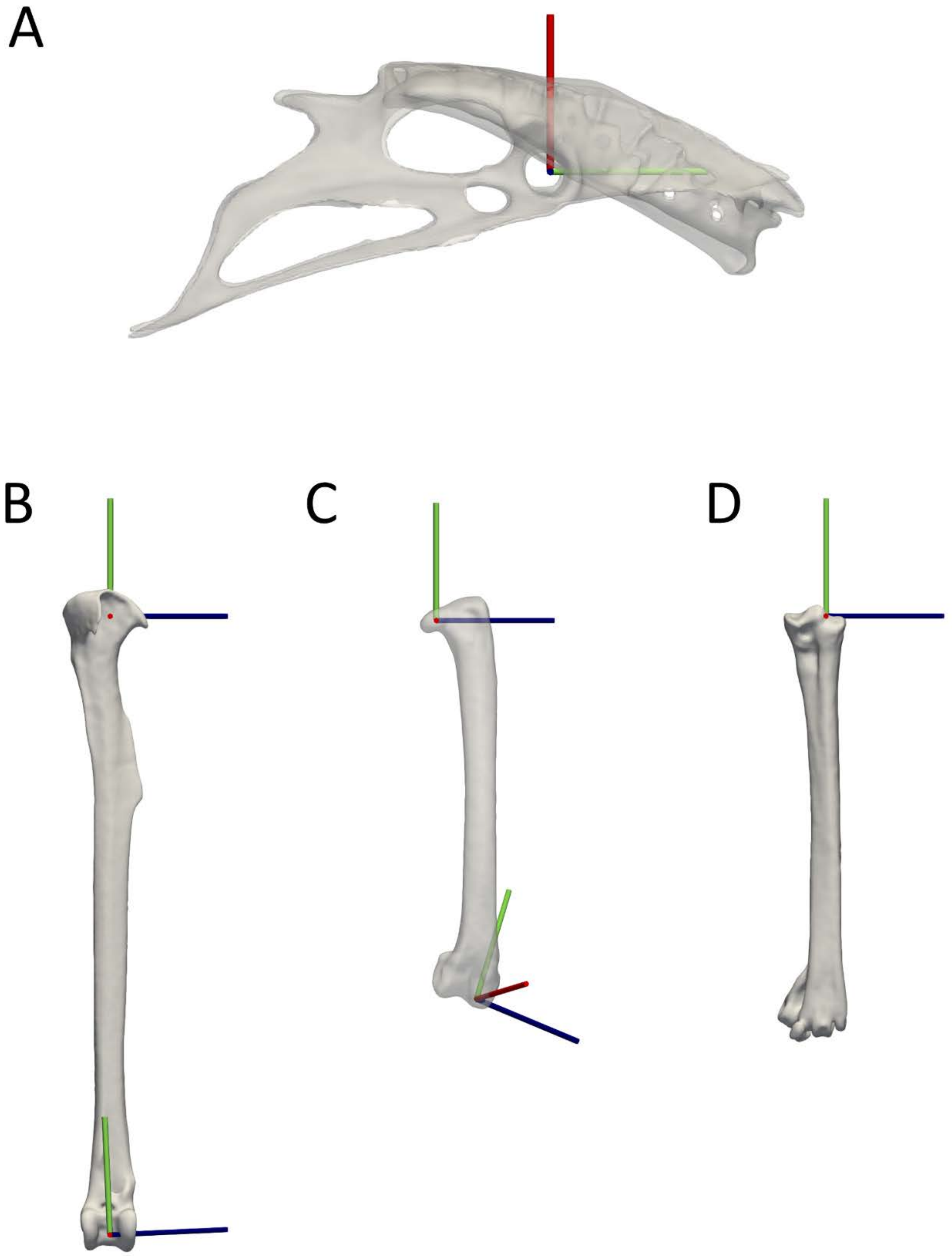


Figure SF5 Anatomical and joint coordinate systems of the left hindlimb bones of the zebra finch. For the pelvis (A), the positive x axis direction (green) points from caudal to cranial, the positive y axis direction (red) points from ventral to dorsal, and the positive z axis direction (blue) points from right to left. For the long bones (B, C and D), the proximal and distal coordinate systems are the anatomical and joint coordinate systems respectively where the positive x axis direction points from distal to proximal, the positive y axis direction points from posterior to anterior, and the z axis direction points from medial to lateral.
